# Psychosocial stress and central adiposity: A Brazilian study with users of the public health system

**DOI:** 10.1101/317891

**Authors:** Flávia Vitorino Freitas, Wagner Miranda Barbosa, Laíz Aparecida Azevedo Silva, Marianna Junger de Oliveira Garozi, Júlia de Assis Pinheiro, Aline Ribeiro Borçoi, Catarine Lima Conti, Juliana Krüger Arpini, Heberth de Paula, Mayara Mota de Oliveira, Anderson Barros Archanjo, Érika Aparecida Silva de Freitas, Daniela Rodrigues de Oliveira, Elizeu Batista Borloti, Iuri Drumond Louro, Adriana Madeira Alvares-da-Silva

## Abstract

**Objective:** To assess the association between indicators of psychosocial stress and central adiposity in adult users of the Unified Health System (SUS) from Southeast of Brazil.

**Methods:** This cross-sectional study was conducted with 384 adults (20 to 59 years old) from the city of Alegre, Southeastern Brazil. The simple random sample represented the population using the public health system of the municipality. The prevalence of obesity was based on the Body Mass Index, and central adiposity (dependent variable) was measured by waist circumference in centimeters. The independent variables were the following indicators of psychosocial stress: food and nutrition insecurity (yes/no), serum cortisol (μg/dL), symptoms suggestive of depression using the Beck Depression Inventory-II ≥ 17 (yes/no), and altered blood pressure ≥ 130/85 mmHg (yes/no). Univariate linear regression was performed between central adiposity and each stress indicator, and later the models were adjusted for socioeconomic, health, and lifestyle variables. All analyses were stratified by rural and urban location.

**Results:** The prevalence of weight excess was 68.3%, and 71.5% of individuals presented an increased risk for metabolic complications related to central adiposity. Mean waist circumference scores for the rural and urban population were 89.3 ± 12.7 cm and 92.9 ± 14.7 cm, respectively (p = 0.012). Indicators of stress that were associated with central adiposity were: cortisol in the rural population and altered blood pressure in the urban population. This occurred both in the raw analysis and in the models adjusted for confounding factors.

**Conclusion:** The associations between stress and adiposity were different between rural (cortisol - inverse association) and urban (altered blood pressure) lifestyles, confirming the influence of local and psychosocial subsistence on the modulation of stress and on how individuals react or restrain stressors. Stress reduction strategies can be useful in public health programs designed to prevent or treat obesity.

## Introduction

Overweight and obesity are defined as abnormal or excessive accumulation of fat that can be detrimental to health.(1) Weight excess affects all regions of the world and is now appearing as a global epidemic. According to the World Health Organization (WHO), 38.9% of the world population aged 18 or more present weight excess and of these, 13.1% are obese.(2) Compared to other WHO regions, the prevalence of weight excess is higher in the Americas (62% for overweight in both sexes, and 26% for obesity in adults over 20 years of age).(3) The prevalence is higher in the United States of America, Mexico, and Chile, where weight excess affects between six and seven out of 10 adults.(3, 4) In Brazil, the National Health Survey (NHS) presented a similar scenario, with 56.9% of overweight adults.(5)

The term stress has already been mentioned in the literature over the years and designates all the non-specific effects of stressors or factors that may act on the body.(6, 7) Stress can be established in the individual in an acute or chronic way, manifested through changes in serum cortisol. Chronic stress can lead to changes in the Hypothalamic-Pituitary-Adrenal (HPA) axis, resulting in altered serum cortisol levels.(8, 9) The frequency of exposure to various stressors can trigger a general adaptation syndrome, composed of three phases: alarm reaction, adaptation phase, and exhaustion phase.(6, 10, 11) The manner in which an individual responds to environmental, economic, social, and health adversities is particularly differentiated, and when the possibilities for adaptation are overcome, psychosocial stress becomes a threat to well-being.(8, 10, 12)

With cultural changes, globalization, and consequent acceleration of the pace of life, there has been an increase in diseases resulting in psychosocial stress and chronic diseases such as obesity, leading concern to public health authorities.(1, 13–19) The relationship between obesity and psychosocial disorders is already well established.(8, 11, 12, 20–22) However, the way different cultures react to the environment and the cause and effect relationship between obesity and stress may be different; these facts along with the issues involved, create a complex concern for authorities. Therefore, this study aimed to evaluate the association between indicators of psychosocial stress and central adiposity in adult users of the Unified Health System (SUS) in a city in the Southeast of Brazil.

## Materials and Methods

### Study design, sampling, and data collection

This was a cross-sectional study with adults (20 to 59 years, SUS users), which was conducted in Alegre, a city located in the Southeast region of Brazil. It is a municipality with the largest population in the micro-region south of the State of Espirito Santo, Brazil, which has a Human Development Index similar to that of the country (HDI = 0.721 and 0.727, respectively), according to the United Nations Development Program (UNDP).(23)

A population of 10222 individuals, registered in the Primary Care Network of Municipal Health was considered in the study design. For the calculation of the simple random sample, absolute accuracy of 5%, 95% confidence interval, design effect equal to 1 and, in the absence of specific studies in the region, it was estimated that the prevalence of overweight individuals was around 50%. Finally, 10% of losses were added and the sample size calculated was 409 individuals.(24)

The study was approved by the Ethics Committee on Human Research, the Health Sciences Center, Federal University of Espirito Santo, under number 1,574,160 issued on June 03, 2016. The inclusion criteria for participation in the study were the following: not being pregnant, having no cognitive conditions that would interfere with the response to the questionnaires, and declare free consent for participation.

Data collection was performed through an individual interview, with questionnaires that evaluated socioeconomic, health and lifestyle conditions, and food and nutrition insecurity (FNiS), as well as symptoms suggestive of depression. Anthropometric, blood pressure, and blood samples were also collected for cortisol analysis.

The socioeconomic, health, and lifestyle questionnaires were elaborated based on the Individual and Domiciliary Registry Files of SUS, Ministry of Health, Brazil. FNiS was evaluated by the application of the Brazilian Food Insecurity Scale (BFIS)(25, 26), a validated instrument comprising 14 questions, aimed at families from the same household with and without members under 18 years of age; concerns evaluated were: lack of food at home, and having some member of the family spending a whole day without eating in the last three months, among others. The degree of severity of FNiS (mild, moderate, and severe) was grouped in this study.

Symptoms suggestive of depression were investigated using the Beck Depression Inventory-II (BDI-II), and total scores were categorized according to the literature: normal, mild, moderate, and severe.(27–29) For the purpose of this study, the following regrouping was used: normal or mild mood disorder (BDI-II <17), and symptoms suggestive of depression (BDI-II ≥ 17).

The anthropometric evaluation was performed by qualified professionals in the morning, after participants had fasted for a minimum of eight hours, following the technical standards of the Food and Nutritional Surveillance System (SISVAN).(30) Stature was evaluated using Alturexata^®^ stadiometer, with a maximum capacity of 2.10 m and accuracy of 0.5 cm. The weight was measured on a Tanita^®^ bipolar bioimpedance balance, with a BC601^®^ branded body fat monitor (with 100 g division and maximum capacity of 150 kg). The Body Mass Index (BMI) was calculated and classified according to WHO reference for adults(1), in which low weight individuals present BMI values < 18.5 kg/m^2^, eutrophic range from 18.5 to 24.9 kg/m^2^, overweight from 25.0 to 29.9 kg/m^2^, and obese individuals BMI being ≥ 30.0 kg/m^2^. Waist circumference (WC) was measured using an inextensible anthropometric tape TBW^®^ (1.5 m and 0.1 cm accuracy), with reference to the midpoint between the last rib and the iliac crest. (31, 32) This study considered that increased risk of metabolic complications for men was associated with abdominal obesity with WC values ≥ 94 cm; and very high risk with WC ≥ 102. For women, an increased risk of metabolic complications was considered at/with WC values ≥ 80 cm, and very high risk, with WC values ≥ 88 cm.(1, 32)

Blood pressure was measured using an aneroid sphygmomanometer with a G-TECH^®^ Premium model. Both the measurement technique and the blood pressure classification were based on the VI Brazilian Guidelines for Hypertension from the Brazilian Society of Cardiology (SBC) for individuals 18 years and older(33), considering altered blood pressure (AP) when values are ≥ 130/85 mmHg.

Blood collection for cortisol analysis was performed in the morning, between 7:00 and 9:00 a.m. The samples were collected in a separator gel tube, kept at room temperature until the clot was retracted, then centrifuged at 2500 rpm for 10 min and refrigerated at 2 ° C to 8 ° C until analysis. The concentration of cortisol was quantified by the chemiluminescence method and the reference values for the morning dose were 6.7 to 22.6 μg / dL.(34)

### Data analysis

Data were tabulated and submitted for consistency analysis. Individuals who presented inconsistency of anthropometric data and those who reported the use of corticosteroid medications were excluded from the analysis.

For the descriptive analysis, the variables were presented as medians and interquartile ranges, or means and standard deviations (non-parametric or parametric data, respectively, according to Kolmogorov-Smirnov normality test), besides frequencies and proportions. The Mann-Whitney U test or Student’s t-test and the chi-square test were used to verify the differences related to the rural and urban locations, considering a level of significance of 5%. In the chi-square test, for the variables that had three or more categories and that presented significant differences, 2 × 2 ratio analysis was performed later, using the Bonferroni correction, which changes the level of significance (p), to avoid type I errors derived from multiple comparisons. After this procedure, the corrected significance level was p<0.016.(35)

Univariate linear regression models were used to evaluate the association between stress indicators and central adiposity. All statistical analyses were performed using SPSS^®^ software, version 15.0 for Windows (IBM, Munich, Germany).

The univariate linear regression analysis consisted of simple models (crude analysis) hierarchically adjusted in three different models: Model 1, analysis adjusted by socioeconomic variables; Model 2, adjusted by socioeconomic and health variables; and Model 3, adjusted by socioeconomic, health, and lifestyle variables.

### Outcome variable: Central adiposity

In this study, the WC in centimeters was chosen as an outcome variable to predict central adiposity from the explanatory variables; it was included in the models as a continuous variable.

### Independent variables

The independent variables selected for the univariate analyses were: FNiS (yes/no), serum cortisol (μg/dL), symptoms suggestive of depression with BDI-II ≥ 17 (yes/no), and AP ≥ 130/85 mmHg (yes/no), which are directly or indirectly related to psychosocial stress. All these indicators were measured by methods and/or validated instruments with good reproducibility in previous studies.

### Potential confounding variables

Socioeconomic variables (age, sex, schooling, and income classification), health (stress report, one or more comorbidities reported, and self-rated health), and lifestyle (current smoking, current alcohol consumption, and weekly physical or leisure activity) were chosen to adjust the univariate linear regression models based on the knowledge collected from the preexisting literature regarding the relation between psychosocial stress and overweight.(7, 8, 12, 14, 36)

Among the socioeconomic variables, it is highlighted that schooling was evaluated in years of study and income was classified according to the Center for Social Policies of the Getúlio Vargas Foundation, which considers low-income individuals with a per capita income of less than US$ 5 per day.(37) In the case of the variable comorbidities, the response options were diabetes, hypertension, cardiac palpitation, cardiovascular disease, and metabolic syndrome, conditions known to be associated with obesity and physical stress.(8) The practice of physical activity was added to leisure, and, as a reference, it was considered that once a week was the minimum frequency to "relief/escape" from the stressful routine, but not to be seen as the ideal.

### Modeling

Beyond the WC (outcome variable), the variables age and serum cortisol entered the models as continuous variables, and all others as categorical and dichotomic, as shown in Table 1.

**Table 1.** Treatment of variables for modeling

| Variables | Dichotomization |
| --- | --- |
| **Dependent variable** |  |
| WC (cm) | continuous variable |
| **Independent variables / Stress indicators** |  |
| FNiS | yes* / no |
| Serum cortisol (µg/dL) | continuous variable |
| BDI-II $\geq 17$ | yes* / no |
| AP $\geq 130/85$ mmHg | yes* / no |
| **Adjustment variables** |  |
| **Socioeconomic conditions** |  |
| Age | continuous variable |
| Sex | female* / male |
| Years of study | < 9* / $\geq 9$ |
| Income classification (< \$5.00/day) | low income* / non-low income |
| **Health conditions** |  |
| Stress report | yes* / no |
| 1 or + comorbidities reported | yes* / no |
| Self-rated health | regular or bad* / good or very good |
| **Lifestyle** |  |
| Current smoking | yes* / no |
| Current alcohol consumption | yes* / no |
| Physical or leisure activities ($\leq 1$ time/week) | yes* / no |

\* indicates the reference category. WC: waist circumference; FNIIS: Food and Nutritional Insecurity; BDI-II: Beck Depression Inventory-II; AP: Amended Pressure.

The univariate analyses for the rural and urban populations were performed separately, to evaluate the differences in the behavior of the predictor variables regarding the stratification of the models by location. All variables used in the models met the assumption of collinearity. The assumptions of normality, linearity, homoscedasticity, and independence of residues were met in all models that had p <0.05 in the F test. Results are presented as non-standard regression coefficients (β), and their respective 95% confidence intervals (CI) and p-values related to the explanatory variable. All analyses were performed using the complete sample, and the significance level was 5% (p <0.05).

## Results

A total of 384 individuals participated in the study, 75 men and 309 women, corresponding to 19.5% and 80.5%, respectively. Among the participants, 133 were rural residents, and 251 lived in urban areas. The median age was 42.5 years (Interquartile Ranges - IR=18.0), and there was no significant difference between the urban and rural populations (p=0.82) or between men and women (p=0.23).

The prevalence of overweight was 33.1%, and that of obesity was 35.2%. Regarding the risk of metabolic complications related to central adiposity (WC, in cm), 51.6% of the sample presented increased risk and the risk of 19.9% substantially increased. The median BMI and WC mean scores were higher in the urban than in the rural populations. The other characteristics related to health, socioeconomic conditions, and lifestyle of the population are presented in Table 2.

**Table 2.** Characteristics of the population according to rural and urban locations

| Characteristics | All | Rural | Urban | p* |
| --- | --- | --- | --- | --- |
| **Health profile** |  |  |  |  |
| **Nutritional Status - BMI** |  |  |  |  |
| Low weight | 1.6 (6) | 1.5 (2) | 1.6 (4) | 0.116** |
| Eutrophy | 30.2 (116) | 33.1 (44) | 28.7 (72) |  |
| Overweight | 33.1 (127) | 36.8 (49) | 31.1 (78) |  |
| Obesity | 35.2 (135) | 28.6 (38) | 38.6 (97) |  |
| **BMI (kg/m<sup>2</sup>) – median (IR)** | **27.5 (7.8)** | **26.8 (7.3)** | **28.4 (8.1)** | **0.027** |
| **Risk metabolic complications - WC** |  |  |  |  |
| Low | 28.5 (109) | 31.8 (42) | 26.8 (67) | 0.084** |
| Increased | 19.9 (76) | 24.2 (32) | 17.6 (44) |  |
| Substantially increased | 51.6 (197) | 43.9 (58) | 55.6 (139) |  |
| **WC (cm) - mean (±SD)** | **91.7 (±14.1)** | **89.3 (±12.7)** | **92.9 (±14.7)** | **0.012** |
| **Classification of cortisol - % (n)** |  |  |  |  |
| Low | 10,0 (37) | 15,0 (19) | 7,4 (18) | 0,042** |
| Normal | 87,8 (324) | 81,9 (104) | 90,9 (220) |  |
| High | 2,2 (8) | 3,1 (4) | 1,7 (4) |  |
| **Serum cortisol (µg/dL) - median (IR)** | **12.1 (6.1)** | **10.8 (5.5)** | **12.9 (6.2)** | **0.006** |
| **Depression - % (n)** |  |  |  |  |
| BDI-II < 17 | 76.4 (272) | 77.5 (93) | 75.8 (179) | 0.728 |
| BDI-II ≥ 17 | 23.6 (84) | 22.5 (27) | 24.2 (57) |  |
| **AP ≥ 130/85 mmHg - % (n)** |  |  |  |  |
| No | 79.8 (296) | 87.5 (112) | 75.7 (184) | **0.007** |
| Yes | 20.2 (75) | 12.5 (16) | 24.3 (59) |  |
| **Stress report - % (n)** |  |  |  |  |
| No | 23.2 (89) | 16.7 (22) | 26.6 (67) | **0.029** |
| Yes | 76.8 (295) | 83.3 (110) | 73.4 (185) |  |
| **Report of comorbidities - % (n)** |  |  |  |  |
| No report | 65.7 (251) | 70.2 (92) | 63.3 (159) | 0.179 |
| One or more comorbidity | 34.3 (131) | 29.8 (39) | 36.7 (92) |  |
| **Use of continuous medication - % (n)** |  |  |  |  |
| No | 45.1 (173) | 46.6 (62) | 44.2 (111) | 0.630 |
| Yes | 54.9 (211) | 53.4 (71) | 55.8 (140) |  |
| **Self-rated health - % (n)** |  |  |  |  |
| Good or very good | 46.9 (180) | 42.1 (56) | 49.4 (124) | 0.173 |
| Regular or poor | 53.1 (204) | 57.9 (77) | 50.6 (127) |  |
| **Socioeconomic profile** |  |  |  |  |
| **BFIS Classification - % (n)** |  |  |  |  |
| FNS | 58.2 (223) | 60.6 (80) | 57.0 (143) | 0.493 |
| FNiS | 41.8 (160) | 39.4 (52) | 43.0 (108) |  |
| **Age (years) - median (IR)** | 42,5 (18,0) | 42,0 (18,0) | 43,0 (18,0) | 0,820 |
| **Skin color - % (n)** |  |  |  |  |
| White | 54.7 (205) | 57.3 (75) | 53.3 (130) | 0.461 |
| Non-white | 45.3 (170) | 42.7 (56) | 46.7 (114) |  |
| **Marital status - % (n)** |  |  |  |  |
| Single | 24.7 (95) | 25.6 (34) | 24.3 (61) | 0.819 |
| Not single | 75.3 (289) | 74.4 (99) | 75.7 (190) |  |
| **Years of study - % (n)** |  |  |  |  |
| ≥ 9 years | 54.0 (204) | 40.9 (54) | 61.0 (150) | **<0.001** |
| < 9 years | 46.0 (174) | 59.1 (78) | 39.0 (96) |  |
| **Income classification - % (n)** |  |  |  |  |
| Non-low income (≥ \$5.00/ day) | 58.5 (224) | 45.1 (60) | 65.6 (164) | **<0.001** |
| Low income (< \$5.00/ day) | 41.5 (159) | 54.9 (73) | 34.4 (86) | |
| **Lifestyle** |  |  |  |  |
| **Physical or leisure activities - % (n)** |  |  |  |  |
| ≥ 1 time/ week | 86.1 (322) | 84.9 (107) | 86.7 (215) | 0.640 |
| Non or ≤ 1 time/ week | 13.9 (52) | 15.1 (19) | 13.3 (33) |  |
| **Current smoking - % (n)** |  |  |  |  |
| No | 91.9 (352) | 94.7 (125) | 90.4 (227) | 0.146 |
| Yes | 8.1 (31) | 5.3 (7) | 9.6 (24) |  |
| **Current alcohol consumption - % (n)** |  |  |  |  |
| No | 68.9 (264) | 78.8 (104) | 63.7 (160) | **0.003** |
| Yes | 31.1 (119) | 21.2 (28) | 36.3 (91) |  |

BMI: Body Mass Index; WC: Waist Circumference; AP: Amended Pressure; BFIS: Brazilian Food Insecurity Scale; FNS: Food and Nutrition Security; FNiS: Food and Nutrition Insecurity; BDI-II: Beck Depression Inventory-II. Quantitative variables presented in medians and interquartile ranges (IR), or means and standard deviations ( ± SD) according to normality (Kolmogorov-Smirnov test); categorical variables presented in relative (%) and absolute (n) frequencies. \* p-value for the tests: Mann-Whitney U or Student t and chi-square, at 5% significance and, when three or more categories, corrected by Bonferroni. \*\* 5% significance, adjusted by Bonferroni ( $p < 0.016$ ).

Tables 3 and 4 present the results of the association between indicators of psychosocial stress and central adiposity in the rural and urban populations, respectively. In the rural population, serum cortisol was inversely associated with central adiposity in all models. AP was associated with adiposity only in the crude model, losing significance in the adjusted models (Table 3). The evaluation done in the urban population showed that only AP was associated with central adiposity and this occurred in all models (Table 4). The other stress indicators evaluated – FNiS and BDI-II> 17 – had no association with central adiposity in the study population (Tables 3 and 4).

**Table 3.** Association between indicators of stress and central adiposity in the rural population

| Stress indicators | Univariate linear regression |  |  | Univariate linear regression models adjusted for confounding variables |  |  |  |  |  |  |  |  |
| --- | --- | --- | --- | --- | --- | --- | --- | --- | --- | --- | --- | --- |
|  | Gross analysis |  |  | Model 1 |  |  | Model 2 |  |  | Model 3 |  |  |
| | $\beta$ | 95%CI | p | $\beta$ | 95%CI | p | $\beta$ | 95%CI | p | $\beta$ | 95%CI | p |
| FNiS | -0.13 | -4.62; | 0.956 | -0.42 | -5.21; | 0.864 | -1.28 | -6.21; | 0.608 | -1.11 | -6.34; | 0.675 |
|  |  | 4.37 |  |  | 4.38 |  |  | 3.65 |  |  | 4.12 |  |
| Serum cortisol | -0.61 | -1.06; | **0.010** | -0.51 | -0.98; | **0.033** | -0.58 | -1.05; | **0.015** | -0.60 | -1.09; | **0.018** |
|  |  | -0.15 |  |  | -0.04 |  |  | -0.11 |  |  | -0.11 |  |
| BDI-II ≥ 17 | 4.62 | -0.71; | 0.089 | 5.16 | -0.06; | 0.053 | 4.31 | -1.45; | 0.141 | 4.95 | -1.57; | 0.135 |
|  |  | 9.96 |  |  | 10.38 |  |  | 10.06 |  |  | 11.47 |  |
| AP | 6.77 | 0.22; | **0.043** | 2.68 | -4.35; | 0.452 | -1.72 | -9.80; | 0.674 | -4.71 | -13.64; | 0.298 |
|  |  | 13.33 |  |  | 9.71 |  |  | 6.36 |  |  | 4.22 |  |

Complex sample. Univariate linear regression. Significance level of 5% (p < 0.05). Dependent variable: Waist circumference (WC) in cm; Stress indicators (independent variables): Serum cortisol (in µg / dL); Food and Nutrition Insecurity – FNiS (yes); Beck Depression Inventory-II – BDI-II ≥ 17 (yes); Amended Pressure – AP ≥ 130/85 mmHg (yes). Gross analysis and univariate linear regression models hierarchically adjusted for confounding factors: *Model 1* - gross analysis adjusted for socioeconomic variables (sex, age in years, schooling, and income status); *Model 2* – gross analysis, adjusted for socioeconomic variables (sex, age in years, schooling, and income status), and health variables (stress report, reporting one or more comorbidities, and self-rated health); *Model 3* – gross analysis, adjusted for socioeconomic variables (sex, age, schooling, and income status), health variables (stress report, reporting of one or more comorbidities, and self-rated health), and lifestyle variables (current smoker, current alcohol consumption, and practice of physical activity and/or weekly leisure). β: beta coefficient; 95% CI: 95% confidence interval; p: p-value. The variables used met the assumption of collinearity. The assumptions of normality, linearity, homoscedasticity, and independence of residues were met in all models that had p < 0.05 by F test.

**Table 4.** Association between indicators of stress and central adiposity in the urban population

| Stress indicators | Univariate linear regression |  |  | Univariate linear regression models adjusted for confounding variables |  |  |  |  |  |  |  |  |
| --- | --- | --- | --- | --- | --- | --- | --- | --- | --- | --- | --- | --- |
|  | Gross analysis |  |  | *Model 1* |  |  | *Model 2* |  |  | *Model 3* |  |  |
|  | β | 95%CI | p | β | 95%CI | p | β | 95%CI | p | β | 95%CI | p |
| FNiS | -0.66 | -4.37; | 0.725 | -0.62 | -4.59; | 0.758 | -1.22 | -5.31; | 0.558 | -0.65 | -4.81; | 0.759 |
|  |  | 3.04 |  |  | 3.35 |  |  | 2.87 |  |  | 3.51 |  |
| Serum cortisol | -0.21 | -0.62; | 0.294 | -0.17 | -0.59; | 0.409 | -0.19 | -0.61; | 0.359 | -0.27 | -0.69; | 0.211 |
|  |  | 0.19 |  |  | 0.24 |  |  | 0.22 |  |  | 0.15 |  |
| BDI-II ≥ 17 | 1.80 | -2.66; | 0.427 | 1.59 | -3.08; | 0.503 | 0.82 | -4.02; | 0.740 | 1.64 | -3.27; | 0.512 |
|  |  | 6.25 |  |  | 6.26 |  |  | 5.65 |  |  | 6.55 |  |
| AP | 9.66 | 5.52; | **<0.001** | 8.41 | -4.05; | **<0.001** | 7.07 | 2.59; | **0.002** | 6.66 | 2.14; | **0.004** |
|  |  | 13.80 |  |  | 12.76 |  |  | 11.55 |  |  | 11.18 |  |

Complex sample. Univariate linear regression. Significance level of 5% ( $p < 0.05$ ). Dependent variable: Waist circumference (WC) in cm; Stress indicators (independent variables): Serum cortisol (in $\mu\text{g} / \text{dL}$ ); Food and Nutrition Insecurity – FNIIS (yes); Beck Depression Inventory-II – BDI-II $\geq 17$ (yes); Amended Pressure – AP $\geq 130/85$ mmHg (yes). Gross analysis and univariate linear regression models hierarchically adjusted for confounding factors: *Model 1* – gross analysis adjusted for socioeconomic variables (sex, age in years, schooling, and income status); *Model 2* – gross analysis, adjusted for socioeconomic variables (sex, age in years, schooling, and income status), and health variables (stress report, reporting one or more comorbidities, and self-rated health); *Model 3* – gross analysis, adjusted for socioeconomic variables (sex, age, schooling, and income status), health variables (stress report, reporting one or more comorbidities, and self-rated health), and lifestyle variables (current smoker, current alcohol consumption, and practice of physical activity and/or weekly leisure). $\beta$ : beta coefficient; 95% CI: 95% confidence interval; p: p-value. The variables used met the assumption of collinearity. The assumptions of normality, linearity, homoscedasticity, and independence of residues were met in all models that had $p < 0.05$ by F test.

## Discussion

This study evaluated the association between indicators of psychosocial stress and central adiposity in adults of a city in Southeast Brazil, and users of the Unified Health System (SUS). The health conditions of the Brazilian population, as well as other nations, transcend the concept of absence of diseases resulting from innumerable social, economic, environmental, and cultural factors. The main health problems of people living in poverty include increased exposure to environmental risk factors, diseases (especially non-communicable diseases), poor nutritional status, difficult access to health services and medicines, as well as other economic, psychosocial, cultural, or health factors, which act as stressing agents, affecting well-being and social interaction.(38)

This study was performed on a population that frequently uses the public health system, which presented socioeconomic (mainly in the rural area) and health fragility, including self-perception reports (53.1% reported poor or regular health), which may represent an impairment of the individual and collective state of well-being, expressing as continuous stress. This, in turn, tends to feed the same cycle of psychosocial instability and lack of health and well-being in a chronic way.

The prevalence of weight excess (BMI ≥ 25.0 kg / m2) found in the present study (68.3%) was equivalent to the current US data and surpassed the weight excess data of the world population (52.0%)(2), of the Brazilian adult population (56.9%)(5), the population of the state of Espirito Santo (52.4%)(39), and of its capital Vitoria (49.7%).(40)

Data from the Family Expense Research - POF 2008-2009(39) showed a lower prevalence of overweight and obesity in rural Brazil (38.8% and 8.8%, respectively), whereas in the present study the prevalence of overweight and obesity were high and statistically similar, both in rural and urban areas (Table 2).

Added to this context, the results were worse when the WC assessment was expanded, with a prevalence of 71.5% at risk of metabolic complications associated with central adiposity, with 51.6% of the individuals at high risk. This proportion exceeded the national prevalence stated in the National Health Survey – NHS(5), which was 37.0%, using the same cut points.(1, 32)

Based on the literature, the main socioeconomic, health, and lifestyle indicators related to excess weight were considered in the present study (Table 2)(5, 9, 14, 26, 39, 41), which have also been referred to in studies as psychosocial stressors.(7–9, 11, 42)

Socioeconomic variables such as schooling and income are indicators of poverty and, together with the FNiS indicator, may represent a direct relation of hunger, and scarcity, with deficiency and malnutrition states.(25, 43, 44) The socioeconomic profile traced in this study pointed to a high prevalence of FNiS (41.8%) in the general population, characterized by low schooling and income, especially in the rural population (Table 2). The nutritional assessment pointed to 30.2% having eutrophy, and only 1.6% having low weight, distancing the population from the limit of 5% that would characterize the presence of current malnutrition.(5) On the other hand, it is known that poverty indicators are also related to excess caloric intake(45) and, consequently, to adiposity. Under these conditions, adiposity may even mask situations of nutritional deficiencies, which is often the case in less developed regions with populations of low schooling and income.(13, 14, 45)

FNiS was one of the variables chosen to evaluate the association between psychosocial stress and adiposity, since this indicator represents scarcity and is related to fear of hunger, insecurity, fragility, and individual and familial instability, which could trigger the activation of the HPA axis, with chronic stress setting. However, FNiS had no association with central adiposity (Tables 3 and 4).

In the context of analysis of health variables, 15% of the individuals living in rural areas had low levels of serum cortisol. Although the prevalence of hypocortisolemia was statistically similar between rural and urban areas, the median serum cortisol was significantly lower in the rural population than in the urban population (p = 0.006), according to Table 2. In addition, in evaluating the association of stress indicators with central adiposity, serum cortisol was inversely associated with central adiposity in the rural population in all univariate analyses (gross analysis and confounding-adjusted models, as in Table 3). In the urban population, serum cortisol was not associated with adiposity (Table 4). The association found in the rural population was contrary to what has been observed in some studies on the relationship between stress and obesity, which have justified the weight gain, due to the increase in intake of palatable and high caloric foods, as a compensatory mechanism resulting in hypercortisolemia of chronic HPA axis activation(7, 8, 20, 46); or weight gain due to the deregulation of satiety mechanisms, through alterations in the leptin system, and also resulting from high levels of cortisol.(7, 12)

On the other hand, the notion that chronic stress acts simply by the direct effect of elevated serum cortisol is becoming less likely relevant in describing how the target tissues respond to cortisol, rather than the levels of the hormone itself.(9) Stressors trigger an HPA axis response by regulating corticotrophin-releasing factor and cortisol feedback and, over time, chronic overexposure to stressors may result in a blunted stress response with decreased levels of cortisol and other associated neurobiological dysfunctions.(47–49) In the absence of supportive care, stressors experienced during life, especially in early life and developmentally sensitive periods, such as pregnancy, childhood, and adolescence, may leave impressions on the neural substrate of emotional and cognitive processes, which may result in a blunted response to cortisol and consequent resilience of the individual to psychosocial stressors.(49) Regarding obesity, hyperactivation of the HPA axis has been observed in obese individuals, but also in individuals with paradoxically normal or low plasma cortisol levels. In a study by research colleagues(50), obesity was associated with relative insensitivity to glucocorticoid feedback. The authors suggested that this condition is characterized by a decreased response of the mineralocorticoid receptor to circulating cortisol.

However, the associations between cortisol and obesity appear to be more complex than anticipated(51). Although it promotes changes in the compensatory mechanism of food intake(7, 8, 20, 46), in the leptin system and satiety control mechanism(7, 8), cortisol is clearly not the only peripheral trigger of adverse effects, which may explain the controversies in this area. It seems likely that the pattern in which cortisol is secreted in front of the stressors is as important as total cortisol secretion. Thus, there is relevance in exploring the social, psychological conditions, health, and lifestyle of a population to identify the main psychosocial stressors and how individuals react and resist to stress.

Another indicator of psychosocial stress used in the present study was depression using BDI-II scores ≥ 17. It is known that early life stress is a risk factor for psychopathology in adulthood, to the extent that the HPA axis was described as deregulated in individuals who experienced early psychosocial stress as well as in those with a wide range of psychiatric disorders.(52) In studies done by colleagues(53) related to chronic stress, depression, and location of urban neighborhoods, the daily stress of living in neighborhoods where residential mobility and material deprivation prevailed was associated with depression. Another study showing the association between urban versus rural environments and depression in the Scottish population found a higher prevalence of psychotropic medication prescription for anxiety, depression, and psychosis in the urban population.(54)

In the present study, a prevalence of depression was found in 23.6% of the population assisted by SUS, with a statistical similarity between rural and urban areas (Table 2). This indicator was not associated with adiposity in any of the models, according to Tables 3 and 4.

In the urban population, the evaluation of health variables pointed to higher levels within the following indicators: median BMI and cortisol, mean WC, and prevalence of high blood pressure (24.3% of individuals with AP), corroborating data from the literature related to hypertension and obesity in urban areas.(55, 56) Lifestyle assessment also showed higher alcohol consumption in the urban population (Table 2). Finally, AP was the only indicator of stress associated with adiposity in the urban population.

It is known that among the main risk factors for elevated blood pressure in adults are obesity, high sodium intake, smoking, alcohol consumption, psychological factors, and certain personality traits such as stress and anxiety.(57–59) Most studies report obesity as a predictor of arterial hypertension(58–63); however, in this study, it was hypothesized that AP, as a psychosocial stress indicator, could predict the increase of WC, which occurred in all models tested (Table 4). It is recalled here that, in the rural population (Table 3), AP was also associated with central adiposity, but this association occurred only in the gross analysis since the adjusted models for confounding variables lost their significance.

Finally, as part of the initial hypothesis, significant differences were observed between home locations. Socioeconomic conditions were challenging in the rural area regarding income and schooling, despite the statistical similarity in the prevalence of FNiS between rural and urban areas. In addition, the report of stress as a personality trait was higher in the population living in the rural region which, on the other hand, presented the lowest median for serum cortisol. As mentioned before, the urban population exceeded the rural population regarding the prevalence of AP. Both the literature review data(53, 54, 64, 65) and the findings of this study reinforce the influence of socioeconomic and livelihood conditions on the different mechanisms of adaptation of the organism to stressors, and on the different forms of coping and developing resilience.

## Conclusions

The stress indicators that were associated with central adiposity were serum cortisol (with an inverse association) in the rural environment and altered blood pressure in the urban area. The study considered several variables known to be associated with stress and weight gain and the models presented robustness in the explanation of the results found.

The observed differences in adiposity prediction regarding housing location reinforce the influence of the local and psychosocial environments on the modulation of stress and on how individuals react to or restrain stressors. Stress reduction strategies can be useful in public health programs designed to prevent or treat obesity.

The study suggests the need for a better evaluation of the use of cortisol as a marker, since both high and low cortisol may explain variations in health status. Both levels relate to different forms of coping or resilience to psychosocial stress. Additionally, owing to the possible bidirectionality of the association between stress and adiposity, new paths should be drawn in the context of a thorough investigation of the mechanisms that explain psychosocial stress, with hypocortisolemia as a biomarker and adiposity as a consequence of current lifestyle.

## Acknowledgments

We would like to thank the volunteers of this study, the Community Health Agents, and the entire Primary Care team of the Municipality of Alegre, ES, Brazil.

## Author Contributions

Conceived and designed the experiments: AMAS FVF. Performed the experiments: FVF WMB LAAS MJOG JAP ARB CLC JKA HP MMO ABA EASF. Analyzed the data: FVF AMAS WMB EBB IDL DRO. Contributed reagents/materials/analysis tools: FVF WMB LAAS MJOG JAP ARB CLC JKA HP ABA DRO EBB IDL AMAS. Wrote the paper: FVF AMAS. Research Project Coordinator: AMAS.

